# Identification of an Aggregation Pheromone from the Small Hive Beetle (Coleoptera: Nitidulidae)

**DOI:** 10.1101/2020.03.03.974741

**Authors:** Charles J. Stuhl, Peter E. A. Teal

## Abstract

Newly emerged adult small hive beetle *Aethina tumida* (Coleoptera: Nitidulidae) emerge from the soil and seek refuge in honey bee hives. Observations of wild and colony reared populations indicate that the beetles form aggregations of many individuals of both sexes. Volatile collections performed on males and females have identified a male produced aggregation pheromone comprised of 6-methyl-5-hepten-2-one, nonanal and decanal. Synergistic effects of the pheromone and a blend of fruit volatiles provide for an effective attractant for both sexes of the small hive beetle. Laboratory assays were performed with the pheromone blend and kairomone blend tested individually combined. This was done using a synthetic aggregation blend along with a fruit-based attractant containing ethanol, ethyl butyrate, acetic acid, ethyl acetate and acetaldehyde. Our results showed that the synthetic aggregation blend along with a fruit-based attractant captured significantly more beetles than the control. The key to a good trapping system is and effective attractant. Our pheromone/kairomone based attractant shows promise to be used as an effective outside the hive control measure for small hive beetle. The identification of the aggregation pheromone is an important step in the search to provide effective control and monitoring of the small hive beetle.

## Introduction

The small hive beetle, *Aethina tumida* Murray is a European honey bee (*Apis mellifera* L, Hymenoptera: Apidae) pest that is destructive to honey bee colonies. Small hive beetle originated in sub-Saharan Africa where it is considered a minor bee pest. Its presence was confirmed in a commercial apiary in Florida in 1998, although previously unidentified specimens indicate its presence in the U.S. since 1996 [1]. Small hive beetle has been a pest on the Australian continent since 2000 [2]. Some species in the family Nitidulidae are found in ripe fruits and melons and are pests of fruit and stored foods [3,4]. Adult beetles and larvae feed on honey, pollen and bee brood [5]. A beetle population can grow exponentially within a short amount of time. Each female can produce ~2,000 eggs in her lifetime and living for many months [6]. A small number of beetles can become tens of thousands of adults in a few months. The larval stage causes severe damage to honey bee colonies resulting in the colony collapsing. Larvae inflict the most destruction by consuming honey bee eggs, brood, pollen and honey. Adults and larvae infect honey and pollen stores with *Kodamaea ohmeri* yeast that causes the honey to ferment, froth and leak from the cells, rendering the honey unsuitable for consumption [7]. When the beetle larval population reaches a certain point, the queen will stop egg-laying and the honey bee colony may abscond from the hive. Weak and queenless hives are more susceptible to small hive beetle damage; however, all colonies are vulnerable to damage when large numbers of beetles are present. Before pupation, the beetle larvae leave the hive and pupate in the soil within proximity to the hives. The larvae remain 3-6 weeks in the soil to complete pupation. Upon emergence, adult beetles will seek refuge in a host bee hive by utilizing colony odors. The adults are strong fliers and easily disperse among hives. Adults can live many months with overlapping generations within in a colony in a single season [8]. The rapid spread of this pest and its impact on the honey bee population has warranted and effective trapping system to reduce its impact on pollination, honey production and honey bee survival. Trapping must be targeted at the adult beetles. Although it is the larvae that cause the damage inside the hive, there is no effective control measure for eliminating them. Therefore, control must target the number of adult beetles in the hive. There have been numerous attempts at developing control strategies for larvae and adults. Honey bee and hive produced volatiles and pollen dough have been used in the past to attract adult beetles [9,10]. Previous studies have utilized corrugated cardboard and corrugated plastic sheets that had been treated with Coumaphos [11]. GardStar® in an insecticidal soil drench applied outside of the hive. This targets the beetle larvae as they pupate in the soil and emerging adults [12]. Unfortunately, none of these control measures are standard practice.

Observations of wild and colony reared populations indicate that the beetles form aggregations of many individuals of both sexes [13]. This observation led to the discovery of a male produced aggregation pheromone. The importance of male produced aggregation pheromones has been well documented from the family Nitidulidae [14,15] and male produced aggregation pheromones have been identified in other Coleoptera such as *Gnathotrichus sulcatus* [16]. In this study we identified the male produced aggregation pheromone to be a combination of 6-methyl-5-hepten-2-one, nonanal and decanal (2:2:1). Small hive beetles have also been shown to be attracted to fruit volatiles, for example cantaloupe has been found to be very attractive to nitidulid and small hive beetles [17,18]. 1992). The main components of the fruit volatile blend have been identified as ethanol, ethyl butyrate, acetic acid, ethyl acetate and acetaldehyde and have shown promise as an attractant for the small hive beetle. Here we describe the investigation of attraction to synthetic version of the pheromone alone and when combined with the cantaloupe volatiles-based attractant. Assays were performed in a wind tunnel and in an environmental chamber using a modified stink bug trap. The target strategy of this system is directed at attraction and capturing small hive beetle adults upon emergence from the soil before they enter the hive. It may also be developed into an in-hive baited trap.

## Materials and Methods

### Source of beetles

*Aethina tumida* were collected from wild populations and then reared in laboratory colonies for two generations. Beetles were collected from honey bee hives maintained at the USDA-ARS, Center for Medical, Agricultural and Veterinary Entomology (USDA-ARS, CMAVE), Gainesville, Florida, USA. All beetles were reared on pollen dough (Global Patties, Butte, Montana) inoculated with *Kodamaea ohmeri* yeast [7]. Beetles were sexed as pupae and placed in moistened soil in separate containers. Insects were reared in a temperature-controlled chamber at 23± 5°C, 60% RH, and photoperiod of 12:12 (L:D) h.

### Volatile Collection

Volatiles were collected separately from 100 adult males and 100 females ~1 week after emergence. All *A. tumida* related collections were performed at the USDA-ARS, Center for Medical, Agricultural and Veterinary Entomology, Gainesville, Florida (CMAVE). Volatiles were collected from male and female beetles using a head space collection technique [19]. Insects were placed in a glass volatile collection chamber (34 cm long and 4 cm outside diameter) with a glass frit inlet and a glass joint outlet and a single port collector base. Collection chamber was covered with a dark cloth and insects allowed to aggregate for 1 h before each collection. Dry charcoal filtered air was pushed into one end the chamber and over the beetles and exited the chamber via a vacuum system. The air then passed through a volatile collection filter containing 50 mg of Tenax^®^ Porous Polymer Adsorbent (Sigma-Aldrich, USA) for 5 min. There were 5 replicates for each sex performed. The isolation of attractive fruit compounds was performed in our lab. This research is out for publication at the same time as writing this manuscript. The fruit volatiles were collected in the same manner as described above.

### Identification of Aggregation Pheromone

The volatile compounds collected were analyzed by GC-MS [GC: Agilent 6890 with an 30 m long HP-5MS capillary column with, 0.25 mm inner diameter, and 0.25-µm film thickness; MS: Agilent 5973 mass selective detector, 70 eV, equipped with an inhouse designed thermal desorption cold trap injector [20] Headspace volatiles collected on Tenax ® TA were desorbed at 220 °C for 2 min by an increased flow of carrier gas (He). The desorbed compounds were trapped and focused by a thermal gradient on the first 5 cm of the column at –78 °C. The separation was initiated by turning of the coolant and allowing the trap to reach the oven temperature by convection heating thus avoiding thermal degradation. The oven temperature of the GC was programmed to rise from 30 °C (3-min hold) to 260 °C at 10 °C/min. The headspace volatiles were identified by comparison of mass spectra with mass spectra libraries [21] and with mass spectra and retention times of authentic standards.

### Electrophysiology Response to Aggregation Pheromone

To determine if male and female small hive beetle had a sensory response to specific compounds isolated from small hive beetle adult males, dilutions of synthetic compounds (Sigma-Aldrich) were exposed to the beetle’s antennae using an electroantennographic detector (EAD). A synthetic blend was created comprised of a 2:2:1 ratio of 6-methyl-5-hepten-2-one, nonanal and decanal. Extracts were analyzed with a GC, split flow interfaced to both flame ionization (FID) and electroantennograph detectors. In this manner, antennal responses were matched with FID signals for compounds eluting from the GC. Volatile extracts were prepared using the above collection set up but with collection on Porapac Q adsorbent and with the collection time extended to 2 h were after the filters were extracted with 150ul of dichloromethane. One μl aliquots were analyzed on a Hewlett-Packard (HP) 5890 Series II gas chromatograph equipped with an HP-5 column (30 m×0.32 mm ID× 0.25 mm) (Agilent, Palo Alto, CA, USA). The oven temperature was held at 40 °C for 5 min, then programmed to increase to 10°C /min to 220°C and held at this temperature for 5 min. Helium was used as a carrier gas at a flow rate of 2.0 ml/min. A charcoal filtered humidified air stream was delivered over the antenna is at 1 ml/min. small hive beetle antennae were excised by grasping the scape at its base with a jeweler’s forceps (No. 5, Miltex Instrument Company Inc, Switzerland). The extreme distal and proximal ends of the antennae were placed in conductivity gel (Parker labs, Fairfield, NJ) between a forked electrode (Syntech, Germany). The electroantennal detector (EAD) and FID signals were concurrently recorded with a GC-EAD program (Syntech EAGPro, Germany), which analyzed the amplified signals on a personal computer.

### Flight Tunnel Bioassay

A flight tunnel bioassay was developed to determine the response of small hive beetle to the synthetic aggregation blend along with a fruit blend. Males, females and a both sexes combined were assayed. There were 10 replicates of each treatment (aggregation pheromone blend, fruit blend, aggregation pheromone/fruit blend). The aggregation pheromone dilution was chosen based on the results obtained from the electrophysiological response to the aggregation blend. The fruit blend was previously developed in our lab from ripe fruit and has shown to be very attractive to small hive beetle. The fruit attractant contained ethanol, ethyl butyrate, acetic acid, ethyl acetate and acetaldehyde. The flight tunnel [22] was constructed of clear acrylic sheets and measured 128 x 31.8 x 31.8 cm and located inside a walk-in environmental chamber at the CMAVE, Gainesville, Florida, USA. Illumination was provided by fluorescent bulbs above the flight tunnel. The light source and the light emitted by the room lighting produced an illumination within the tunnel of ~1600 lux. The room temperature ranged from 28.7-28.8 º C and humidity between 37.6 −38.1% RH. Air flow within the tunnel was produced by a Shaded Pole Blower (Dayton, Niles, IL) which pulled air into the tunnel through a charcoal filter and exhausted it outside the chamber. The exhaust end was screened to prevent insects from entering the tube. Airflow could be adjusted using a baffle inside a tube that connected the downwind end of the tunnel with the exhaust system of the hood. Air speed was maintained at 0.2 m/s. This flow was determined to be the speed that most stimulated flight in small hive beetle.

Two 3.8 L glass jars fitted with a metal lid containing two brass hose fittings contained the fruit and allowed air to pass over the odor source and the blank control and emerge separately in the flight tunnel. Air flow into the fruit containers was controlled by an adjustable flow meter (Aalborg Instruments, Monsey, NY) set at ~0.5 LPM Treated air emerged into two insect traps located at the upwind end of the tunnel and placed midway between its ceiling and floor. These were constructed from 40-dram clear plastic snap cap vials (Thornton Plastics, Salt Lake City, UT). A 10 mm hole was placed in the center of the cap to allow insects to enter the chamber. Twenty-five males and twenty-five females were placed in the flight tunnel and was checked every 0.5 h from the period of 0900 to 1400 hours. A positive response was recorded when there was a beetle inside the trap. The insect was removed from the trap and replaced with a naive insect from a stock cage where the original insects had been obtained. The position of the treatment and control were changed after each replication to prevent positional effects. There were 10 replicates performed for each concentration blend. Concentrations were selected from GC-MS results and quantified using known standards.

### Trapping bioassay

Trapping assays were performed in a climate-controlled chamber at 23± 5°C, 60% RH, and photoperiod of 12:12 (L:D) h. An inverted Rescue Reusable Stink bug trap (Sterling International, Inc., Spokane, WA) was used in the assay. The trap was inverted to allow for the entrance to face upright. Two traps, a treatment and a blank control were suspended from the ceiling of the enclosure. The treatment contained the aggregation pheromone + fruit blend. This was presented in the same manner as the flight tunnel assay. The attractant blend was delivered via an impregnated 3 cm cotton dental place inside a 1 ml Eppendorf® tube (Sigma-Aldrich, USA). The open tube was then attached to the inside chamber of the trap. A blank control was run alongside the treatment, with the treatments being ~1 meter apart. A vial containing 200 male and 200 female newly emerged small hive beetle adults was opened inside a screen mesh cylindrical field cage (91.5 c diam. x 183 c height). The assay was run for 24h after which the trapped insects were counted. The position of the treatment and control were changed after each replication to prevent positional effects. There were 10 replicates performed.

### Statistical analysis

Analysis of data was performed using ANOVA [23]. When variables proved to have an insignificant effect on numbers of males and females captured, data were pooled and pair-wise comparisons of responses to treated and control fruit accomplished with the Wilcoxon paired-sample test [24].

## Results

### Identification of Aggregation Pheromone

Three components were isolated from male small hive beetle. The most abundant component was 6-methyl-5-hepten-2-one and two minor components, nonanal and decanal (Fig 1a). The identical compounds were also isolated from small hive beetle females, but in a very small amount compared to what was being produced by the males.

**Figure 1.**
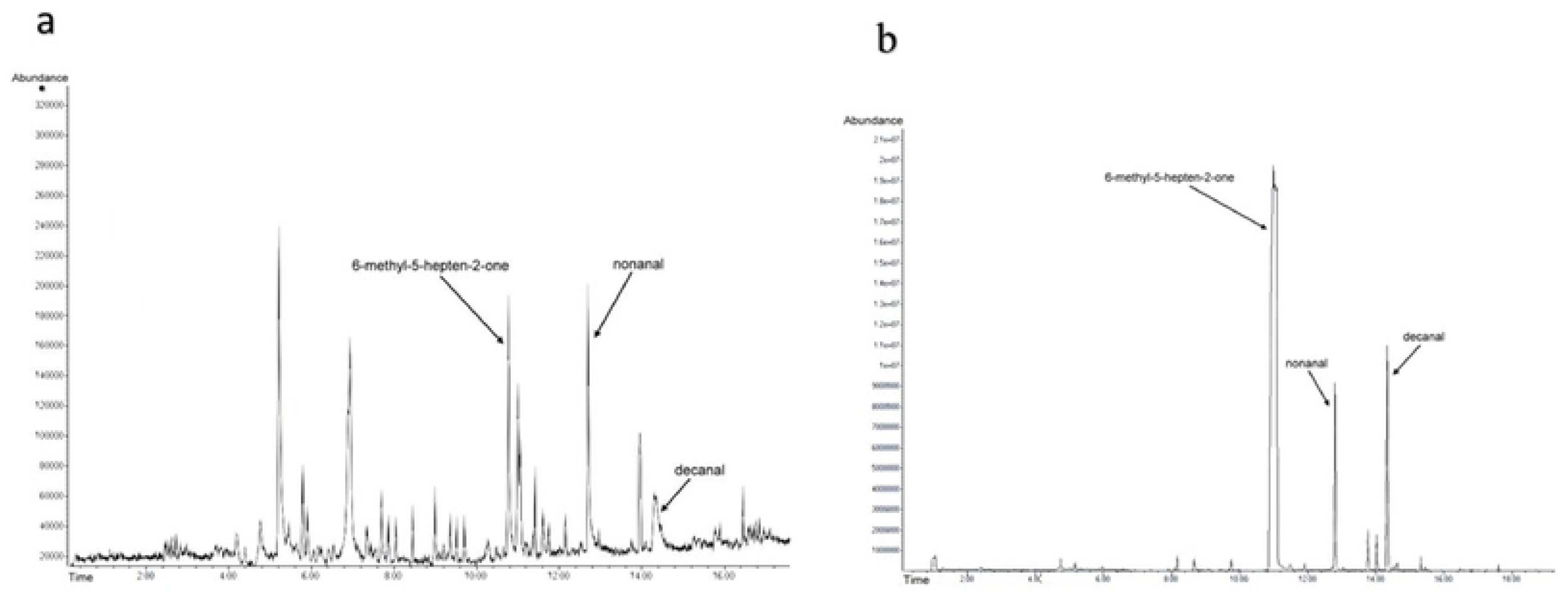
a) GC-MS chromatogram of *A. tumida* aggregation pheromone; b) GC-MS chromatogram of *A. tumida* synthetic aggregation pheromone

### Electrophysiology Response to Aggregation Pheromone

Sensillae on the antennae of small hive beetle males and females responded to the natural and synthetic aggregation blend. The greatest response was to 6-methyl-5-hepten-2-one, nonanal and decanal. This procedure allowed for the evaluation and selection of the active compounds that initiated an electrophysiological response (Fig. 1b).

### Flight Tunnel Bioassay

The flight tunnel assays results indicated a distance attraction to the aggregation pheromone blend containing 6-methyl-5-hepten-2-one, nonanal and decanal (Fig 2a). The fruit blend had a better attraction than the aggregation blend (Fig 2b). There was a significant increase in trap capture when the aggregation blend was used in conjunction with the fruit volatile odor (p>0.0001) (Fig 2c).

**Figure 2.**
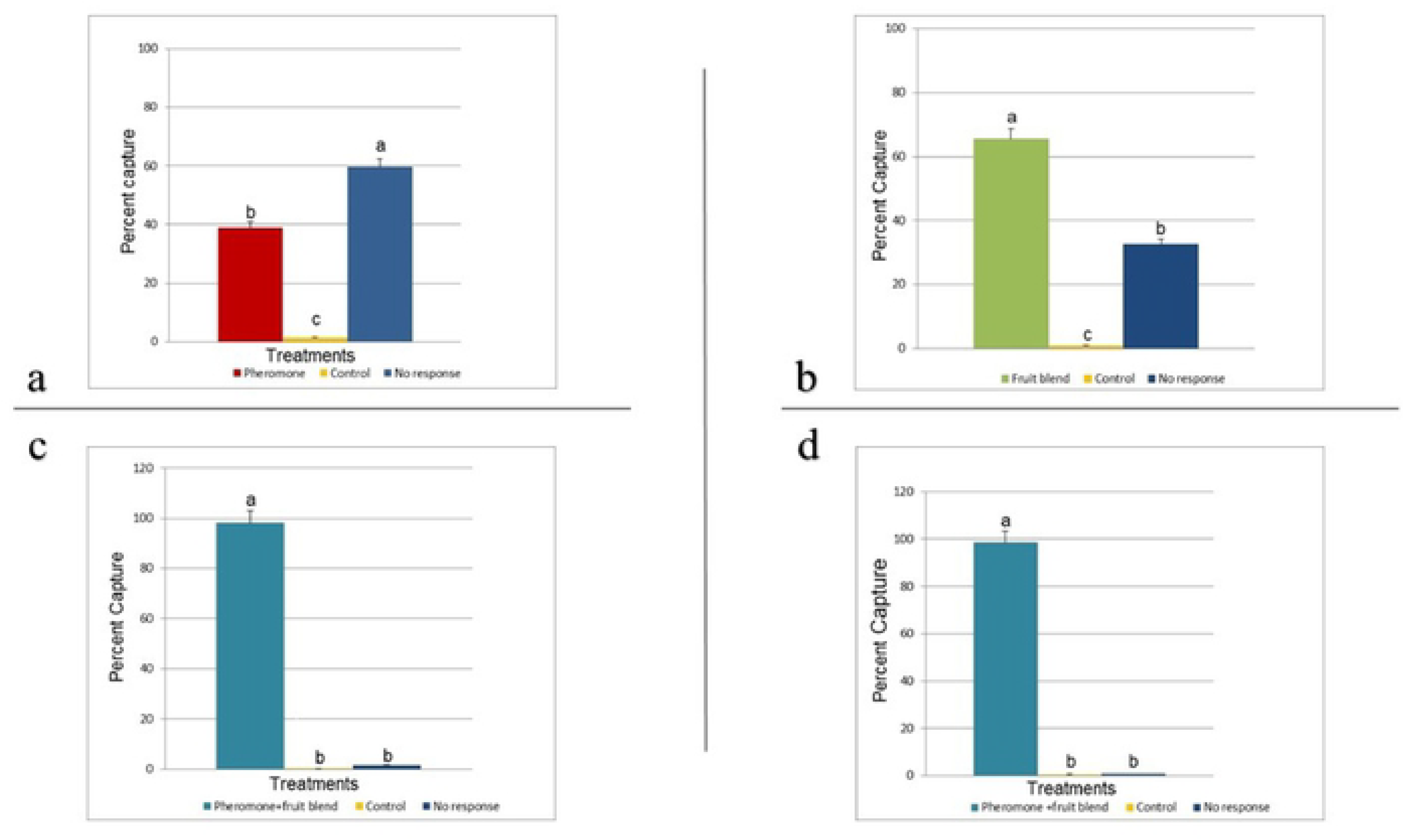
Percent capture of the flight tunnel response of *A. tumida* to a synthetic blend of chemicals that mimic the male produced aggregation pheromone, a) the aggregation pheromone blend was 39%, blank control 1.5%, no response 60% (p>0.0001); b) the fruit volatile blend was 65%, blank control >1%, no response 32.6% (p>0.0001); c) the aggregation pheromone+fruit volatile blend was 98%, blank control >1%, no response >1% (p>0.0001); d) the trap response for the aggregation pheromone+fruit volatile blend was 99%, blank control >1%, no response 1.7% (n=3641, p>0.0001). Percentage values are associated with the various sets of bars. Those sharing a letter are not significantly different.

### Trapping Bioassay

The trapping bioassay confirmed the strong wind tunnel attraction to the combination blend compared to blank controls (p>0.0001) (Fig. 2d).

## Discussion

This study demonstrated that a blend of three compounds isolated from small hive beetle induced a high level of attraction. GC-MS analyses identified the aggregation blend comprising 6-methyl-5-hepten-2-one, nonanal and decanal. These compounds have never been reported as associated with small hive beetle. The pheromone was isolated from male and female (in a much lower amount) small hive beetle and was found to be attractive to both sexes. A synthetic blend of chemicals in a 2:2:1 was formulated into a blend that mimics the small hive beetle aggregation pheromone elicited a positive antennal physiological response. Although the insects tested had an antennal response to individual compounds, only the blend initiated a behavioral response as expected for a pheromone.

Although the aggregation pheromones alone showed activity in the wind tunnel assay, the attraction was poor when compared with fruit odors. The lowest amount of captures was seen when the aggregation pheromone was used alone, with a significant number of beetles not responding. When the fruit odor was presented alone it attracted 65% of the released beetles while the paired blend attracted 98% of the beetles. A similar synergistic effect has also been seen for other insects like *Carpophilus lugubris* [25] and the maize weevil [26] The control and beetles that did not respond were <1%. When placed in a trapping device, along with the fruit odor, the pheromone attracted 99% of the released weevils. The control and no response were <1% respectively.

Trap captures indicate that the male produced aggregation pheromone is detected and produces a behavioral response in both sexes of adult beetles including aggregation once collected in the trap.

It is not possible to control this pest within a hive by means of an insecticide without harming the honey bees. A baited trap that that is directed at the small hive beetle and restricts honey bee access would be a very attractive way to target this pest within and outside of the hive. This method has been successfully demonstrated for monitoring and the reduction of nitidulid beetle populations [15]. The most successful integrated pest management eradication of an agriculture pest was the Boll Weevil Eradication Program. The discovery of the male produced aggregation pheromone of the cotton boll weevil, *Anthonomus grandis*, led to the development of an eradication strategy [27]. This system used pheromone traps for weevil detection, cultural practices by modifying the beetles habitat to decrease its food supply, followed by chemical treatments that reduced the beetle’s weevil populations.

An integrated pest management approach for small hive beetle management may be accomplished with the use of the synthetic aggregation pheromone paired with fruit odor blend. Additional research will focus on the possible sex bias and methods to lengthen the pheromone activity over an extended period. This may be accomplished by placing the pheromone/kairomone blend on many available release matrixes. We will further investigate and develop an improved trap that will better contain the insects and prevent escapes. The attractant blend is highly attractive and has the potential to be extremely successful in trapping the small hive beetle. Thus, this discovery has the potential to control an invasive species that is affecting honey bee survival worldwide.

## Acknowledgments

I thank Dr. Jennifer Gillett-Kaufman (University of Florida, Entomology and Nematology Department) and Dr. Hans Alborn (USDA-ARS-CMAVE) for their critical review of this manuscript. The use of trade, firm, or corporation names in this publication is for the information and convenience of the reader. Such use does not constitute an official endorsement or approval by the United States Department of Agriculture or the Agriculture Research Service of any product or service to the exclusion of others that may be suitable.

